# Accounting for central place foraging constraints in habitat selection studies

**DOI:** 10.1101/2022.10.21.513205

**Authors:** Simon Benhamou, Nicolas Courbin

## Abstract

Habitat selection studies contrast the actual space use with the expected use under the null hypothesis of no preference (hereafter neutral use). Neutral use is most often assimilated to the relative abundance of the different habitat types. This generates a considerable bias when studying habitat selection by foragers that perform numerous back and forth to a central place (CP). Indeed, the increased space use close to the CP with respect to distant places reflects a mechanical effect rather than a true preference for the closest habitats. Yet, correctly estimating habitat selection by CP foragers is of paramount importance for a better understanding of their ecology and efficiency of conservation actions. We show that (1) including the distance to the CP as a covariate in unconditional Resource Selection Functions (RSFs), as did in several studies, is quite inefficient to correct for the bias. Bias can be eliminated only by contrasting the actual use distribution to an appropriate neutral distribution that takes the CP forager behavior into account. (2) The need to specify such an appropriate neutral use distribution can be bypassed by relying on a conditional RSF, where the neutral use is assessed locally without reference to the CP.

## Introduction

Since the seminal paper by Johnson (1980), the preferences that animals may have for different types of resources and habitat features has become an important question in theoretical and applied ecological studies (Svanbäck & Bolnick 2005, Fortin et al. 2008, McLoughlin et al. 2010). Addressing such a question about habitat or resource selection makes it possible to identify environmental drivers that shape animal space use and to predict population distributions (Boyce et al. 2002, Manly et al. 2002, Johnson et al. 2013, Fieberg et al. 2021), and thereby to improve conservation and management actions (Thurfjell et al. 2014, Courbin et al. 2022).

A key concept in habitat selection analyses is the selection ratio *w*_*x,y*_ = *u*_*x,y*_/*a*_*x,y*_, where *u*_*x,y*_ and *a*_*x,y*_ are the values at location (*x*, *y*) of the probability density functions (PDF) of actual use and expected use under the null hypothesis (H_0_) of no preference for any particular environmental feature, respectively. One of the most important feature that may drive space use in numerous terrestrial animal species is the local habitat type, i.e. the discrete category of the habitat in which the location (*x*, *y*) occurs (e.g. ‘forest’ and ‘meadow’). The selection ratio for a given habitat type *H*, irrespective of the other features to which an animal may be sensitive, is *w_H_* = *U_H_*/*A_H_*, with *U_H_* = ∬_*H*_ *u*_*x,y*_ *dxdy* and *A_H_* = ∬_*H*_ *a*_*x,y*_ *dxdy*, where ∬_*H*_ *dxdy* means that integration is done for all locations (*x*, *y*) occurring within habitat type *H*.

Classically, *U_H_* and *A_H_* refer to the use and the availability of habitat type *H*, respectively. The latter is often considered a synonym of relative abundance. Following Johnson (1980), Rosenberg & McKelvey (1999) and Matthiopoulos (2003) rightly considered it should rather be understood as a synonym of accessibility. Indeed, the expected use of habitat type *H* under the null hypothesis of no particular preference, *A_H_*, corresponds to the relative abundance of type *H* only when all types are equally accessible. This occurs only if *a*_*x,y*_ is uniform or if the proportions of the various habitat types are not correlated with a spatial feature (e.g. the distance to given location) to which *a*_*x,y*_ may depend on. To avoid any ambiguity, hereafter we will refer to *A_H_* as the ‘neutral use’ of habitat type *H*. Many habitat selection studies focus on the third selection order, i.e. how an animal uses space within its home range (Johnson 1980). In the particular case of central place (CP) foragers, the neutral use PDF, *a*_*x,y*_, cannot be expected to be uniform because of the numerous back and forth to the CP, where the neutral use should peak. Individuals of many taxa behave as CP foragers, either during the major part of their adult life, such as hymenopterans (e.g. Kacelnik et al. 1986), or during the breeding period, such as birds (e.g. Ventura et al. 2022) and wolves (e.g. Ylitalo et al. 2021). Assuming that *a*_*x,y*_ for a CP forager is uniform should therefore result in flawed habitat selection ratios (Appendix S1), except if the habitat types are distributed at random with respect to the CP.

However, over the past decades, habitat selection has been mainly studied with Resource Selection Functions (RSFs, Manly et al. 2002, Johnson et al. 2013, Fieberg et al. 2021). Rosenberg & McKelvey (1999) claimed that, for a CP forager, the selection ratios for the various habitats types could be correctly assessed within the classical (unconditional) RSF framework by comparing the actual use distribution with a neutral use distribution simply reflecting the relative abundances of the different habitat types while including the distance from the CP as a covariate. Although this approach was used in a number of habitat selection studies (e.g., Bond et al. 2009, Chudzińska et al. 2015), we doubted that it could provide reliable selection ratio estimates, because the actual distribution of distances to the CP may be poorly related to the appropriate neutral distribution. In this paper, using toy examples, we investigated to which extent a misspecification of the neutral use of the different habitat types simply reflecting their relative abundances can be corrected by including CP distance-based covariates in unconditional RSF models when applied to CP foragers. Alternatively, selection ratios can be estimated locally in a matched design (local actual use being paired with local resource availability; Compton et al. 2002, McDonald et al. 2006, Duchesne et al. 2010, Prima et al. 2017). As these conditional RSF models should relax the spatial constraint due to the CP, some authors (e.g. Carrete & Donázar 2005, Carlson et al. 2021) assumed that they provided unbiased estimates of habitat selection ratios for CP foragers, but this was not yet demonstrated. Using computer simulations, we checked whether the CP constraint can be efficiently relaxed in this way. Finally, we proposed guidelines to improve studies of animal habitat selection under severe spatial constraints.

## Methods

Because of the unit sum constraint on the (actual and neutral) use of the different habitat types (Σ_*H*_*U*_*H*_ = Σ_*H*_*A*_*H*_ = 1), the selection ratios for the different types have a limited meaning by themselves, as they depend on the set of types that are considered relevant for the analysis (Johnson 1980). In contrast, the ratio of selection ratios for any two habitat types is independent of the arbitrary set composition. It therefore constitutes a reliable metric of the preference for a given habitat type relatively to another, on which we relied to assess the suitability of the different approaches we tested. In any case, we considered a forager exploiting a circular home range from a place located at the center of its home range.

To determine whether, in the unconditional approach, including the distance to the CP as a covariate makes it possible to correct the selection ratio estimates when the neutral use PDF *a*_*x,y*_ is wrongly assumed to be uniform, we contrasted two RSF models. One focused only on the habitat type whereas the other included both the habitat type and the distance *D* to the CP. With habitat type 1 taken as reference, the type-distance RSF model was expressed as:

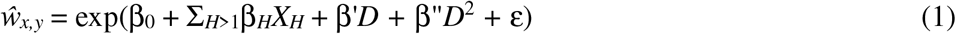

where *X_H_* is an indicator function (often called a dummy variable in the RSF context) set to 1 if location (*x*, *y*), at distance *D* from the CP, lies in habitat type *H* or to 0 otherwise, Σ_*H*>1_ corresponds to a sum over all types *H* > 1, β are the various parameters to be estimated through likelihood maximization, and ε is a random error term. Although *D* was expected to affect 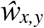 in a monotonic way, we considered a quadratic form to allow a more flexible fit. The type-only RSF model was written as the type-distance model (Equation 1) with β’ and β” set to 0, i.e.

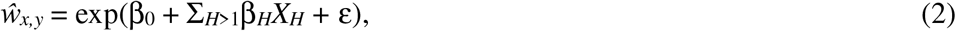

which can be rewritten simply as 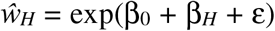 for any type *H* > 1. In both models,the preference for any type *H* > 1 relatively to type 1, *w*_*H*_/*w*_1_, is estimated by exp(β_*H*_). The correct values of β_*H*_ are those obtained with the type-only model based on the approprinaeteutral use. For a CP forager, setting a neutral use in the type-only model that simply reflects the relative abundances of the different habitat types (uniform neutral use PDF) should result in biased β_*H*_ values, except if the relative proportions of habitat types are independent of the distance to the CP. Looking at the exp(β_*H*_) values obtained with the type-distance model based on a uniform neutral use PDF allowed us to determine whether including the distance to the CP can or cannot correct a misspecification of the neutral use.

In this unconditional RSF approach, we contrasted two kinds of neutral use PDF for *D≤R,* where *R* is the home range radius (Fig. 1A). One corresponds to the classical, but misspecified for a CP forager, bivariate uniform law (*a*_*x,y*_ = (π*R*^2^)^−1^). The other corresponds to a bivariate exponential law (*a*_*x,y*_ = 1.5/(πσ^2^) exp(–√3*D*/σ), with σ^2^ = *E*(*D*^2^)/2 = 0.75 *E*^2^(*D*), that is the distribution expected, in the absence of particular habitat preference, for a forager relying on a CP-biased correlated random walk (Benhamou 1989, 1996). We truncated this distribution at the 0.95 cumulative isopleth (*D* = 2.74 σ) and divided the *a*_*x,y*_ values by 0.95 to obtain a cumulative distribution that sums to 1 within *R* of the CP. We also contrasted three kinds of actual use PDF for *D≤R* (Fig. 1A): bivariate uniform (*u*_*x,y*_ = (π*R*^2^)^−1^), conical (*u*_*x,y*_ = 3(1–*D*/*R*)/(π*R*^2^)) or reverse conical (*u*_*x,y*_ = 1.5(*D*/*R*)/(π*R*^2^)), reflecting typical space use behaviors. The conical PDF roughly matches the neutral use PDF, but with a reduced contrast between the highly used nearby areas and the marginally used faraway areas. The reverse conical PDF mimics what happens with some colonial seabirds that avoid exploiting the area close to the colony (continental shelf) and preferentially exploit distant productive areas (e.g. continental slope, oceanic fronts) by commuting between the CP and these distant areas. Finally, the uniform PDF corresponds to an intermediate situation where the forager tends to compensate the mechanical overuse of areas close to the CP (e.g. by using a speed that is inversely related to *D*).

**Figure 1.**
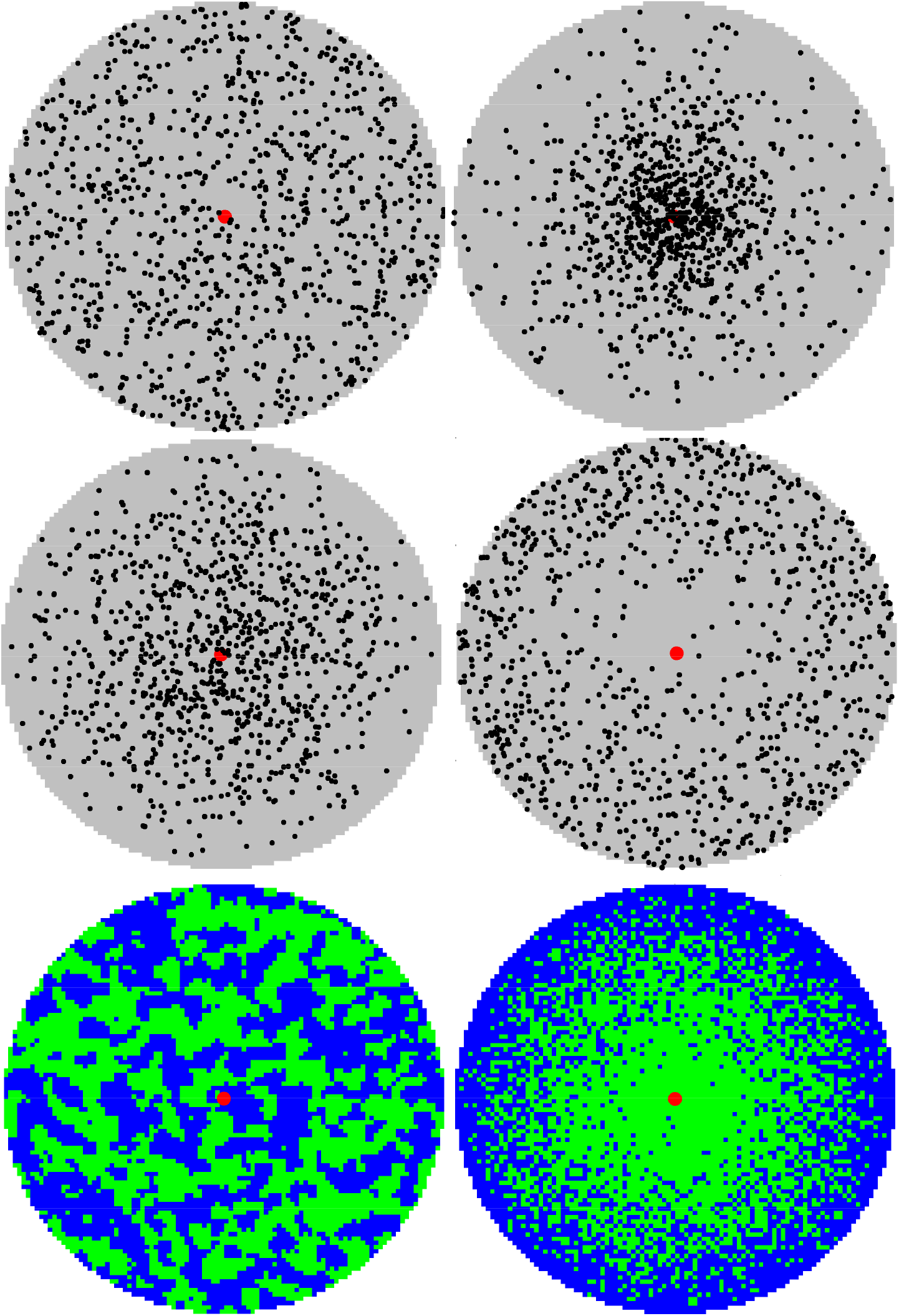
Use distributions and landscape configuration. A) The four types of PDF involved are, from top to down and left to right: bivariate uniform, bivariate exponential, conical and reverse conical. For clarity, only 1000 locations (black dots) are shown in addition to the CP (red dot), whereas the results were obtained by drawing 100,000 locations. B) Two landscape configurations: (left) ‘CP-independent’ landscape, where the proportions of habitat types 1 and 2 are independent of the distance to the CP, and (right) ‘CP-dependent’ landscape, where the proportion of habitat type 2 is proportional to the squared distance to the CP.

For simplicity, but without loss of generality, we modelled the landscape within the home range as made of two equally abundant habitat types. They were mixed according to two landscape configurations. In the ‘CP-independent’ configuration, the local proportions of cells of types 1 and 2 did not depend on the distance to the CP (Fig. 1B). With this configuration, we aimed only to check whether, when the CP cannot bias the preference for one of the two type, the relative preference of habitat type 2 over type 1 is correctly estimated by an unconditional RSF model where the neutral use distribution simply reflects the relative abundance of the two types. In the ‘CP-dependent’ configuration, the local proportion of cells of habitat type 2 ranged from 0 to 1 proportionally to the squared distance to the CP (which warranted that, overall, the two habitat types were equally abundant, and resulted in a correlation between the habitat type and the squared distance of 0.58; Fig. 1B). We took habitat type 1 as the reference, and therefore looked at the value of exp(β_2_) as the relative preference of habitat type 2 over type 1. We drew 100,000 ‘actual’ locations from uniform, conical or reverse conical PDF within the *R*-radius circle, and 100,000 ‘neutral’ locations (usually called pseudo-absences) from exponential or uniform distributions within the same circle. We generated these PDFs as well as the two landscape configuration with the R (v3.6.2, 2019) packages *NLMR* (Sciaini et al. 2018), *raster* (Hijmans 2021) and *sp* (Bivand et al. 2013). Relative preferences were estimated using an unconditional logistic regression (Manly et al. 2002, Keating & Cherry 2004, McDonald 2013), based on the expression 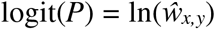 or 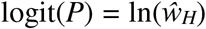, where *P* corresponds to the probability that a location in habitat type *H* at distance *D* from the CP belongs to the whole set of actual rather than the whole set of neutral locations.

Based on the same ‘CP-dependent’ landscape, we determined whether the relative preference of habitat type 2 over type 1 could be suitably estimated for a CP forager within a conditional RSF framework. We simulated a CP-biased correlated random walk made of 100,000 steps with a length equal to the cell size. A walker with no particular preferences was assumed to move a step per unit time. A preference for habitat type 2 cells *k* times larger than for type 1 cells was simulated by designing a walker that stayed *k*–1 unit times without moving each time it reached a type 2 cell. We estimated the relative preference of habitat type 2 over 1 with a conditional logistic regression (R package *survival*, Therneau 2021) that paired the habitat type of the cell actually used at any given unit time with the habitat types of the 8 non-chosen cells out of the 3×3 cells surrounding the animal’s location at the previous unit time.

## Results

As expected, in the ‘CP-independent’ landscape, the relative preference for habitat type 2 over type 1 was correctly estimated to be equal to 1 (i.e. no particular preference for any type) by all unconditional RSF models. This held true whatever the way the actual use PDF depended on the distance to the CP (uniform, conical or reverse conical), even if the neutral use was wrongly assumed to simply reflect the relative abundances (uniform neutral use PDF). In contrast, when the relative proportions of the two habitat types depended on the distance to the CP, with type 1 being the majority at close distance and type 2 at large distance, there was a clear relative preference for type 2 over type 1 for the three types of actual use considered, as shown by the type-only model based on the appropriate (i.e. exponential) neutral use PDF (Table 1A). With an inappropriate (uniform) neutral use PDF, the estimation of this relative preference was biased, resulting in estimates that were more than 4 times lower than the correct values for the three actual use PDF (Table 1A; for a conical actual use, the relative preference of habitat type 2 over 1 was even reversed). Incorporating the distance to the CP in the unconditional RSF resulted in an absence of relative preferences. Thus, the bias was only partly corrected when the actual use PDF was conical, not corrected when the actual use PDF was uniform, and even increased when the actual use PDF was reverse conical (Table 1A). Overall, incorporating the distance to the CP into the unconditional RSF model to hopefully correct the bias due to an inappropriate specification of the neutral use distribution is just hazardous. Specifying the actual use and the neutral in polar rather than Cartesian terms (Appendix S2) and/or including an interaction term between habitat types and distance *D* to the CP (as Rosenberg & McKelvey 1999 did; Appendix S3) did not lead to any improvement.

**Table 1.** Relative preference of habitat type 2 over type 1 (i.e; 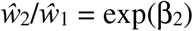) estimated for the ‘CP-dependent’ landscape, which is characterized by an increasing proportion of habitat type 2 with the distance to the CP. (A) Results obtained with the unconditional RSF models; (B) Results obtained with the conditional type-only RSF model. The correct values are written in bold.

| A) Unconditional RSF<br>Model | Neutral use | Actual use |  |  |
| --- | --- | --- | --- | --- |
|  |  | Conical | Uniform | Reverse<br>conical |
| Type-only | Exponential | **1.78** | **4.09** | **6.19** |
| Type-only | Uniform | 0.43 | 0.99 | 1.49 |
| Type-Distance | Uniform | 0.99 | 0.98 | 0.99 |
| B) Conditional RSF | | Simulated relative preference $k$ | | |
|  |  | **1** | **2** | **3** |
| Estimated relative preference |  | 0.98 | 1.97 | 2.97 |

The relative preference of habitat type 2 over type 1 was suitably estimated in the ‘CP-dependent’ landscape using a conditional logistic regression (Table 1B). We also obtained the same results simply by computing the ratio of the mean local selection ratios, with the local actual use being equal to 0 or 1 depending on whether the animal reached a cell of the type considered or of the other type, and the local neutral use based on the relative abundances of the two types (when both present) in the 3×3 cells surrounding the previous location.

## Discussion

We showed that a strong spatial constraint on animal movements, such as the need to come back frequently to a CP, largely affects how the landscape is used but cannot be considered just an additional factor. Spatial constraints should be included as so in unconditional habitat selection approach when designing the neutral use distribution, i.e. the distribution expected under the null hypothesis of no particular preferences for some habitat types over some others. Contrary to the claims of Rosenberg & McKelvey 1999, including this constraint as an additional covariate while designing the neutral use distribution simply as reflecting the relative abundances of the different habitat types is unlikely to provide correct estimates. In contrast, a conditional approach, by which the neutral distribution is estimated based on local resource availability makes it possible to determine the correct values of relative preferences.

A major difficulty for the unconditional RSF approach is to design a suitable neutral use PDF. Fortunately, the simple uniform PDF, which means that the expected time spent in any habitat type simply reflect its relative abundance in terms of area, is a suitable choice when the proportion of the different habitat types is independent to the distance to the CP. The user should therefore always check this assumption beforehand at the population level (for non-colonial CP foragers, the independence may arise at the population level even if, for each individual, there is a strong correlation between the distance to the CP and the proportions of habitat types, when individual trends differed and thus may compensate each other). In the general case where independence cannot be granted, the suitable choice of the neutral use PDF strongly depends on what we know about the way the animal tends to move. Some authors assumed, without sound justification, that the expected density *ax,y* may decrease linearly with distance to the CP (Matthiopoulos 2003, Wakefield et al. 2011). Here we assumed that, in the absence of any preference, the animal performs search loops (from and to the CP) that are driven by a biased correlated random walk with a constant bias strength, which results in a bivariate exponential neutral use PDF (Benhamou 1989, 1994; see Monsarrat et al. 2013 for an application example). However, there are other possibilities. In some species, such as *Cataglyphis* ant (Wehner & Wehner 1990), individuals search for resources during the outward journey and come back to their nest along a straight line once they had caught a prey item. One can guess that such animals rely on a simple (unbiased) Correlated Random Walk, i.e. a purely diffusive movement, during the outward journey, with a diffusion coefficient that depends on the habitat type. In this case, the analysis of habitat selection may focus on the space use distribution during the outward journey with a neutral use PDF corresponding to a bivariate Gaussian distribution centered on the CP and a variance reflecting the mean diffusion coefficient (see Codling et al. 2008). More generally, a neutral use distribution can be obtained by simulating movements starting from and ending at the CP independently of the habitat features (Aarts et al. 2008, Raymond et al. 2015, Hazen et al. 2016, Courbin et al. 2018, 2022). For animals that commute between their nest and a distant habitat (e.g. as occurs with breeding pelagic seabirds that exploit oceanic fronts, e.g., Wakefield et al. 2011, Kappes et al. 2015, Courbin et al. 2018), one may rely on a uniform neutral use PDF after having removed the commuting part, which strongly reflects the CP spatial constraint.

Interestingly, suitable habitat selection ratio estimates were obtained with a conditional RSF approach (Compton et al. 2002, McDonald et al. 2006, Duchesne et al. 2010, Prima et al. 2017), which bypasses the need to explicitly specify the whole neutral distribution. Indeed, in conditional approach (including more sophisticated forms such as Step Selection Function models; Fortin et al. 2005, Thurfjell et al. 2014, Avgar et al. 2016), the neutral use is assessed only locally from the local resource availability. A conditional habitat selection analysis (e.g. Carrete & Donázar 2005, Carlson et al. 2021), makes it possible to relax the spatial constraint due to the CP that results in places that are all the less accessible as they are far away from the CP (Rosenberg & McKelvey 1999, Matthiopoulos 2003). Such an approach may sometimes problematic because it rests on the assumption that the process of habitat selection depends only on the local conditions. However, in the present context, ignoring the larger scale movement processes is clearly an advantage. In the simple case where only habitat types are considered, the relative preferences can also be suitably estimated as the ratio of the mean local selection ratios.

A related question is whether distance to the CP should be included as a covariate in a habitat selection analysis. When the proportion of habitat type is not correlated to this distance, unbiased results about preferences for the different habitat types can be obtained with a uniform neutral PDF. In contrast, when such a correlation occurs, this may be problematic. With an unconditional RSF model, it is preferable not to include the distance in the model because its effect is assumed to be accounted for by a neutral use PDF assumed to be appropriate (usually without guarantees). This issue can be avoided by relying on the conditional approach, but this will not necessarily result in additional insight. For example, in our study where the proportion of the habitat type 2 is proportional to the squared distance to the CP, including the distance (in simple or quadratic form) is useless (i.e. the relative preference in terms of habitat types remained unchanged and the selection coefficient(s) associated to the distance were almost null). Furthermore, in both unconditional and conditional approaches, the distance and habitat type becomes mutually confounding factors, so that results should be interpreted with caution.

Other strong spatial constraints may affect space use, in particular the occurrence of areas that are hardly accessible. For example, for colonial pelagic birds that nest on a continental cliff and systematically avoid flying above the ground, space use cannot be expected to be evenly distributed in all directions around the nest. Similarly, in territorial species where individuals actively defend their whole home ranges, the areas belonging to conspecifics’ territories are hardly accessible. Even if they are not defended, remote areas (with respect to the home range core area) tend to be avoided as they are not part of the home range. Such spatial constraints must be taken into account when designing the neutral use PDF (see examples in Wakefield et al. 2011, Raymond et al. 2015, Fluhr et al. 2017, Courbin et al. 2022).

Overall, we demonstrated the importance of properly specify the neutral use distribution in unconditional habitat selection analyses, or to shortcut this specification by relying on a conditional approach. This should be a primary consideration when evaluating habitat selection of CP forager because of the strong potential negative consequences for the understanding of species ecology and the efficiency of conservation actions.

