## Supplementary information for "Accounting for central place foraging constraints in habitat selection studies"

### S1- Relative abundance vs. neutral use

Below is illustrated in 1D the discrepancy that may exist between relative abundances and expected use under the null hypothesis of no preference (i.e. neutral use). When the distribution of habitat types depends on the distance to central place, the ratio of relative abundances of habitat types 1 and 2 (represented by the lengths of the green and the blue bars) can suitably represent the ratio of expected use under the null hypothesis of no preference (neutral use, represented by the green and blue areas) only when the distribution is uniform (top panel). When the distribution is centrally peaked (bottom panel), the neutral use of habitat types that occur close to (distant from) the central place is clearly under- (over-) estimated by its relative abundance.

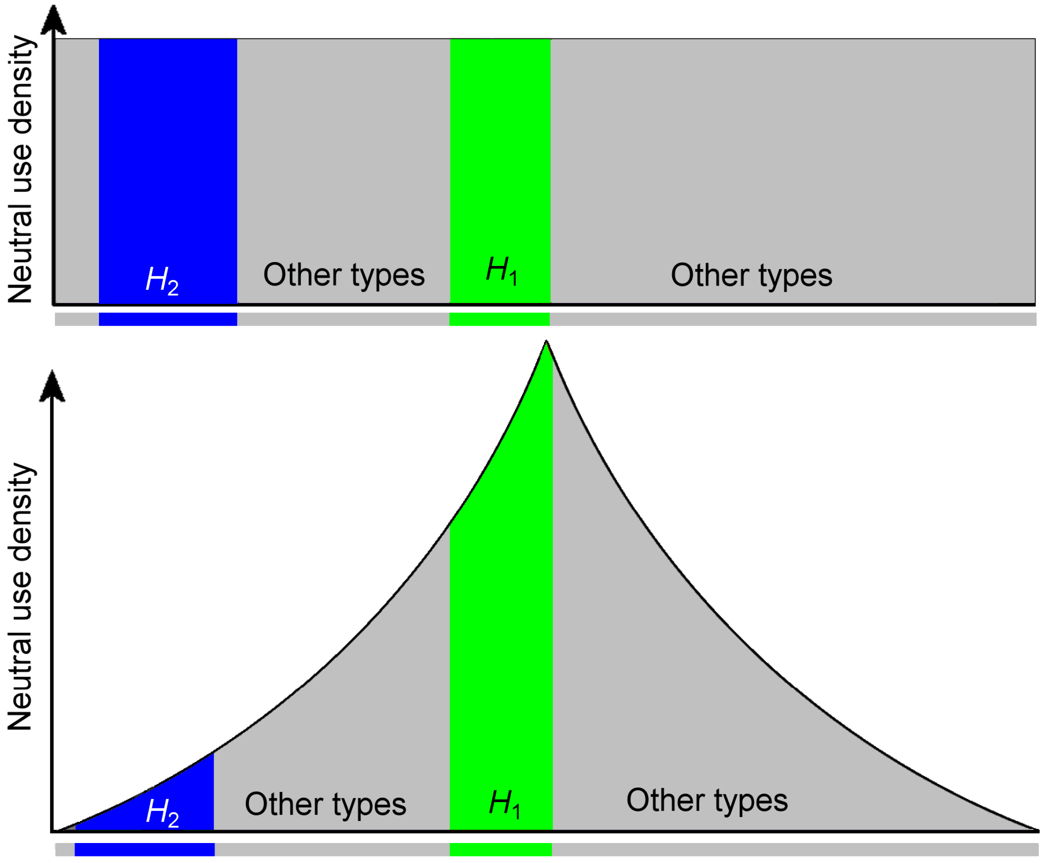

**S2 - Specification of the models in Cartesian vs. polar terms**

One may wonder whether the unconditional RSF model would have been able to correct (at least partially) for the bias if the actual use and neutral use were expressed as functions of the distance $D$ to the central place rather than in Cartesian terms. The corresponding PDF (per unit of arc length) are $u_D = 2\pi D u_{x,y}$ and $a_D = 2\pi D a_{x,y}$, where $u_{x,y}$ and $a_{x,y}$ are actual use and neutral use PDF (per unit area), respectively. Thus, despite the area of a ring with radius $D$ and infinitesimal width d$D$ is proportional to $D$, this dependence on $D$ affects the actual use and the neutral use per unit of arc length in the same way. Consequently, the selection ratios are not affected by the choice of relying on a Cartesian or polar approach.

**S3 - Including interaction terms**

One also may wonder whether the inability of our type-distance model to correct for the bias would result from the absence of an interaction term between habitat types and distance to the central place $D$. We did not include such a term because, as expected, a wrong specification of the neutral use results in a wrong estimate of selection ratios for the different habitat types only when their proportions are correlated with the distance. In such a case, habitat types and distance act at least as partially as confounding factors, making the selection ratios hard to interpret. If selection ratios for the different habitat types are further allowed to be dependent on distance through the inclusion of interaction terms, the results would become fully unintelligible. However, to be sure we did not miss something, we considered the more general RSF model including the interaction term between habitat type and distance (with associated regression coefficient $\beta_{inter}$) $\hat{w}_{x,y} = \exp(\beta_0+\beta_2 X_2+\beta' D+\beta'' D^2+\beta_{inter} X_2 D+\varepsilon)$, with $D$ ranging between 0 and $R$. The distance-dependent ratio of selection ratios for habitat types, $\hat{w}_2/\hat{w}_1 = \exp(\beta_2+\beta_{inter} D)$, remained close to 1 for the three types of actual use (uniform: from 1.05 to 0.94, conic: from 1.29 to 0.85; reverse conic: from 0.94 to 1.01). Thus, even if there was no issue of interpretability, the inclusion of the interaction term does not help reduce the bias resulting from using an inappropriate uniform neutral use PDF.
